# Auditable recovery of single-cell RNA-seq zeros with SPARE

**DOI:** 10.64898/2026.06.02.729664

**Authors:** Sergio Hernández-Galaz, Ignacio Pezoa-Soto, Andrés Hernández-Oliveras, Sofía Rodriguez, Alvaro Lladser, Alberto J.M. Martin

## Abstract

In Single-cell RNA-seq, observed zeroes are the mix between biological absence and technical limitations. However, current evaluation metrics fail to distinguish between these two states, focusing on reconstruction accuracy rather than the biological validity of edits. We introduce SPARE, a partition-aware framework that audits the imputation process by cataloging observed zeros as edited, unchanged, or marker-vetoed prior to sequence reconstruction. Across multiple tissue benchmarks, SPARE successfully recovered masked expression while cataloging edit burden, marker leakage and raw-state movement. Protein and disease-context audits showed that recovered expression must be accepted, restricted or rejected endpoint by endpoint. SPARE reframes imputation as auditable zero editing rather than generic matrix completion.

## 1 Background

In single-cell RNA sequencing (scRNA-seq), computational imputation aims to resolve extreme data sparsity by inferring observed zeroes across individual cells. Traditionally, this process is evaluated as a prediction problem: observed expression values are hidden, methods reconstruct them, and performance is summarized by reconstruction error (Talwar et al. 2018; Batson et al. 2019; Hou et al. 2020; Cheng et al. 2023). This approach, while useful, does not capture the biological intervention made when an imputer changes an observed zero. Once modified, the recovered value can alter marker interpretation, differential-expression rankings, cell-state boundaries and downstream biological claims (Andrews and Hemberg 2018; Kim, Zhou, and Chen 2020; Hou et al. 2020; Cheng et al. 2023). Consequently, downstream interpretation depends not only on how accurately a value is reconstructed, but on which observed zeros are edited at all (Hou et al. 2020; Cheng et al. 2023; Linderman et al. 2022). This distinction also echoes broader missing-data benchmarks, where reconstruction error can diverge from task utility (Zhou, Bouadjenek, and Aryal 2024).

At the core of this challenge lies the context-dependent nature of single-cell sparsity, as observed zeros are inherently heterogeneous. These zeros are composed of both biological absence of gene expression and technical artifacts derived from missed capture, limited sequencing depth or stochastic low expression (Qiu 2020; Hou et al. 2020; Sarkar and Stephens 2021; Jiang et al. 2022; Miao et al. 2025). In the context of UMI-based data, zero patterns often carry cell-type and cell-state information, transforming the significance of a zero into cellular context rather than technical noise (Li and Li 2018; Linderman et al. 2022; Qiu 2020; Jiang et al. 2022; Svensson 2020). Consequently, the interpretation of a zero becomes context-specific. For example, a missing lineage marker may be a plausible candidate for recovery in one cell state, while in another, it represents an essential biological boundary better left unchanged (Kim, Zhou, and Chen 2020; Cheng et al. 2023; Linderman et al. 2022). Thus imputation must not be viewed as a dataset-wide instruction to fill missing values, but as a context-aware decision about editing measured non-detection.

Paradoxically, existing frameworks lack an explicit mechanism for documenting the transition from measured non-detection to reconstructed signal. Current strategies operate as terminal processes where raw data is ingested and a denoised matrix is returned, while the internal rationale for each modification is discarded (Dijk et al. 2018; Hou et al. 2020). This results in the absence of a record of interventions, leaving researchers unable to verify whether a zero is edited, bypassed as signal, or vetoed due to biological risk. Without this auditable trail, reconstruction accuracy, edit burden, and marker-context risk become impossible to separate This is specially problematic when imputed expression is used to support high-stake biological endpoints such as marker discovery, differential expression, rare-state interpretation or disease-context annotation (Andrews and Hemberg 2018; Kim, Zhou, and Chen 2020; Hou et al. 2020; Cheng et al. 2023).

To mitigate these risks, the field has moved away from unconstrained matrix completion towards more biologically and mathematically restricted models. Initial advances focused on structural properties and dropout distributions, utilizing approaches such as ALRA’s zero-preserving low-rank strategy (Linderman et al. 2022) or frameworks like scRecover, PbImpute, and scZiva that explicitly model true-zero versus dropout-zero states (Miao et al. 2025; Y. Zhang et al. 2025b; Shamsuzzaman, Ray, and Mukhopadhyay 2026; Vo et al. 2026; Tran et al. 2022). More recently, methods have incorporated explicit biological safeguards to reduce false activation, with tools such as SmartImpute restricting recovery to panels predefined marker (Yao et al. 2025) and frameworks like SCRABBLE and scZN leveraging external constraints (Peng et al. 2019; Wu et al. 2026). Recognizing further the context-dependent nature of this process, recent approaches utilize cellular partitions to make zero recovery more biologically constrained, this is by summarizing local expression programs, marker constraints, and detection structures via reliable clusters, transferred labels and curated annotation (Y. Zhang et al. 2025a), these partitions identify contexts where recovery is plausible versus those where it would be misleading. Thus cellular partition aware frameworks like CPARI, which combines cell partitioning with imputation-position detection (Y. Zhang et al. 2025a), separate themselves from their label-free counterparts by the integration of cellular context directly into the recovery process.

However, despite these conceptual advances, a fundamental gap remains in how imputation is evaluated and recorded. Benchmark debates have shown that semi-synthetic recovery remains insufficient as methods that improve masked-value reconstruction can still induce false marker signal, distorting downstream analyses, and altering the nonzero fraction, drifting from measured raw-state summaries (Huang et al. 2018; Andrews and Hemberg 2018; Li and Li 2019; Huang and Zhang 2019; Valyaeva et al. 2026).To diagnose these specific distortions, marker-based false-positive leakage has been used to assess whether imputation blurs cell-type-specific expression (Wu et al. 2026), yet, efforts to fully audit this leakage are limited by the outputs of the algorithms themselves. Thus, even context-aware frameworks collapse the measured and synthetic states and obscure their edits. The resulting loss of an intervention record precludes the audit of these algorithmic decisions, meaning researchers can measure downstream distortions but cannot quantify their origins.

Here, we introduce Selective Partition-Aware Recovery of Expression (SPARE), implemented in the Python package spare-sc (imported as spare).By separating the decision to edit an observed zero from the value assigned during reconstruction, the framework catalogues these interventions before values are overwritten. Using a configured group_key to calculate within-partition support, contextual priors, and marker-safety constraints, the algorithm materializes a selected-zero intervention map prior to writing any values. This map explicitly catalogues entries as measured nonzeros, selected observed zeros, untouched observed zeros, and marker-vetoed zeros, carrying this exact provenance forward into recovery, burden, leakage, raw-state, partition-sensitivity, and external-support audits.

By returning this map, evaluation shifts away from a single reconstructed matrix and toward a co-primary audit vector that catalogues downstream effects across four technical metrics: recovery, which tracks the independent selection and reconstruction of masked nonzeros; edit burden, which quantifies the volume of observed-zero space rewritten by the selected-zero mask; marker leakage, which identifies edits occurring in marker-negative, off-label, or externally unsupported contexts; and raw-state movement, which measures the downstream displacement of marker rankings, differential-expression overlap, or raw-state concordance away from the un-imputed state. Under this architecture, burden describes where values are written, whereas raw-state movement catalogues how much the downstream analysis changes after writing. Collectively, these four metrics provide the analytical foundation for evaluating partition-sensitivity and external-support contexts throughout our validation audits.

We evaluate SPARE not as a universal replacement for existing imputers, but as an auditable intervention framework that catalogues how recovered expression can be accepted, restricted, or rejected across diverse experimental and biological conditions. We demonstrate these validation boundaries using matched benchmarks, external marker panels, paired CITE-seq protein measurements, and ovarian-cancer disease-context audits. While SPARE is not the first method to treat single-cell zeros as heterogeneous, preserve zeros, target selected entries, or utilize cellular partitions to guide recovery, its distinct contribution is the inspectable intervention map it returns. By making the zero-editing decision explicit before reconstruction, this catalog provides the necessary tracking unit to systematically evaluate marker safety, leakage, edit burden, and downstream biological interpretation.

## 2 Results

### 2.1 SPARE records zero-editing decisions before imputation

An observed zero should be edited only when the evidence for recovery justifies changing a measured non-detection. SPARE makes that decision explicit by separating zero selection from value reconstruction. For each gene-cell pair, the framework catalogs measured expression, selected observed zeros, marker-vetoed zeros and untouched observed zeros. This intervention map is generated before any reconstructed values are written,preserving it as an auditable output for downstream validation.

The operational logic of the SPARE framework bridges data heterogeneity, zero selection, and downstream auditing (Fig. 1). Within this pipeline andidate observed zeros are eligible for recovery only when statistical, neighborhood and partition-level evidence support expression and no marker-safety veto applies (Fig 1a,b). Thus under this architecture the final output pairs the reconstructed matrix with a co-primary audit trail, explicitly linking each recovered value to its selection criteria, localized cellular context, and quantitative edit burden (Fig. 1c).

**Figure 1.**
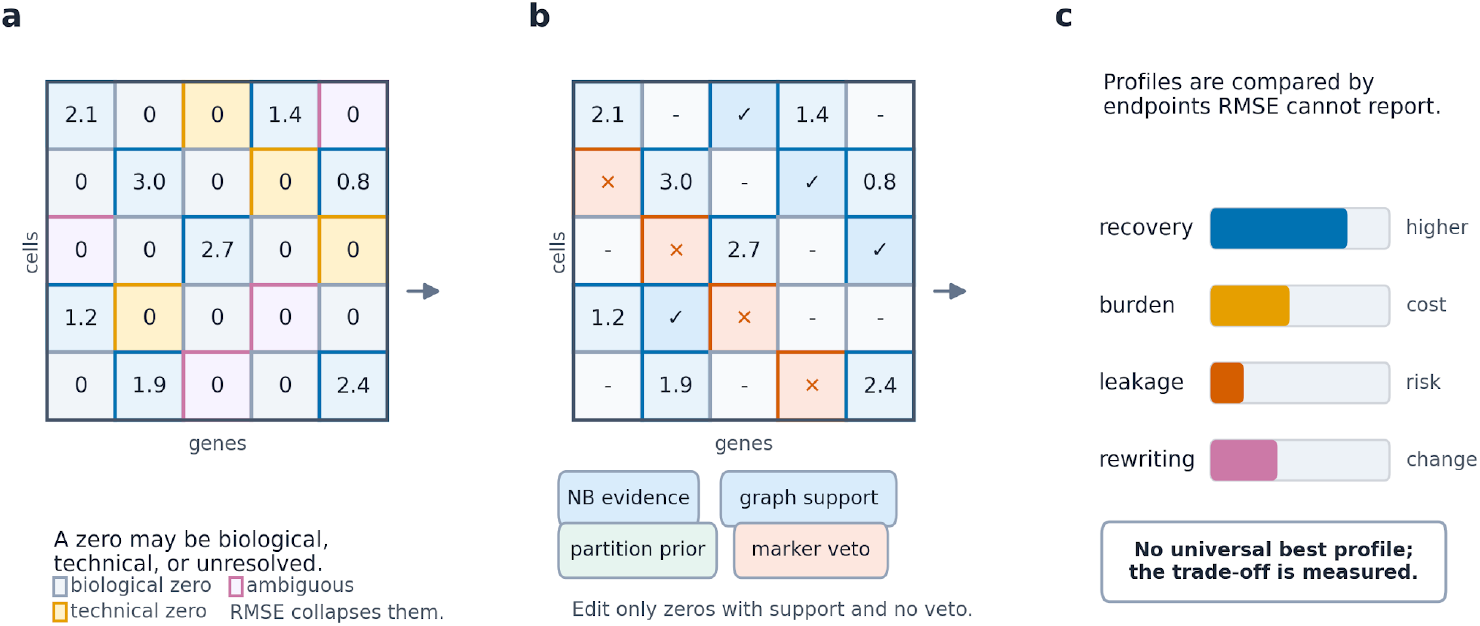
SPARE makes zero recovery an auditable intervention. a, Observed zeros in scRNA-seq mix biological absence, missed capture and limited evidence. b, SPARE separates the decision to edit an observed zero from the value assigned after reconstruction. The selected-zero intervention map distinguishes measured nonzero entries, selected observed zeros, marker-vetoed zeros and untouched zeros before values are written. c, Each recovered value is interpreted through an audit vector that includes recovery, edit burden, marker leakage, raw-state movement and external support. The figure defines the decision framework used in the analyses below.

**Figure 2.**
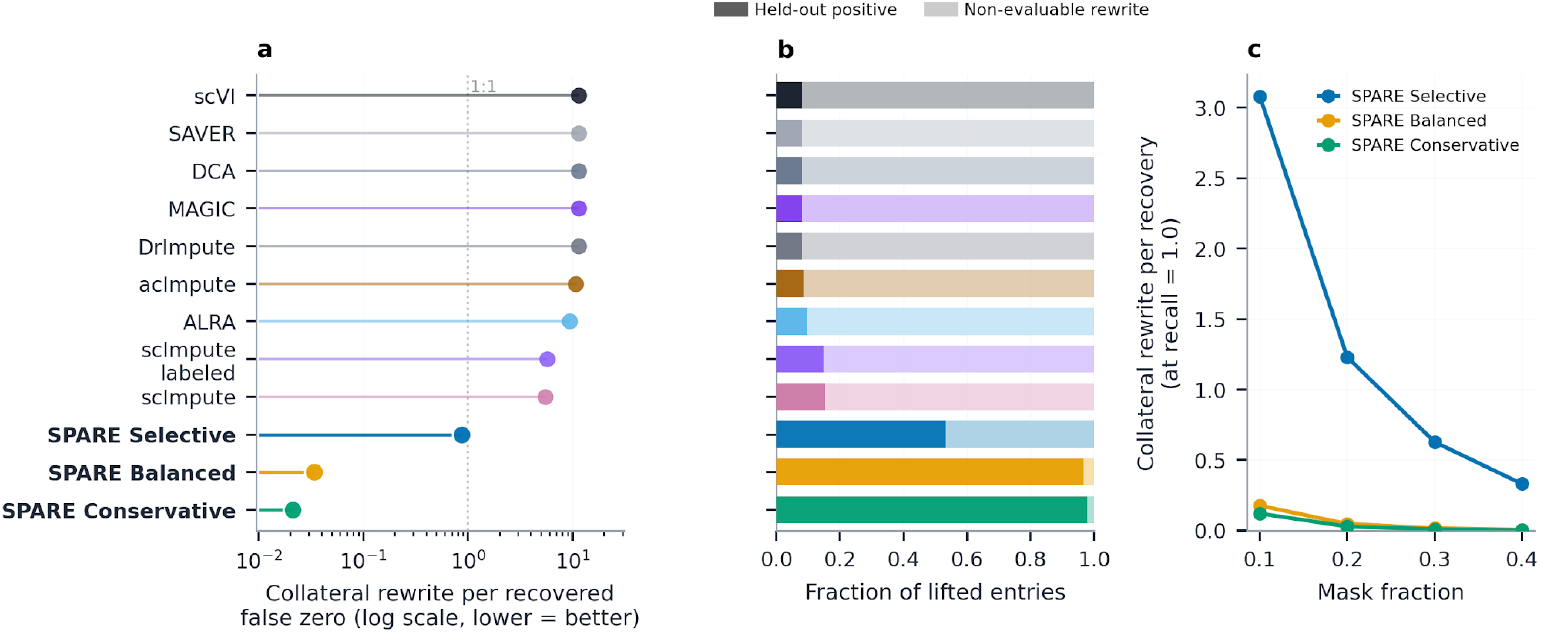
Zero-selection audits expose recovery yield, collateral rewriting and profile sensitivity. a, Collateral rewriting per recovered false zero in the PBMC comparator audit. Values near 1 indicate a roughly one-to-one relation between recovered held-out positives and additional selected observed-zero entries; larger values indicate broader rewriting outside the evaluation target. b, Composition of lifted entries for the same comparator rows. Bars separate held-out positives from non-evaluable rewritten zeros, showing whether apparent recovery is concentrated in the masked target set or accompanied by broader collateral editing. c, SPARE profile sensitivity across mask fractions. Selective, Balanced and Conservative profiles show different collateral-rewrite costs as masking becomes more difficult, making the recovery-cost operating point explicit.

To demonstrate that observed-zero editing is biologically defensible, our validation results systematically evaluate the framework across three core axes (see Fig. 1). We first test leakage-safe recovery of held-out expression, quantify the collateral rewriting required to obtain that recovery, and verify whether edited entries remain compatible with baseline marker rules, raw-state summaries and external biological evidence. These performance benchmarks are formalized into an endpoint-level decision guide (see Table 1) for interpreting recovered expression. Within this framework, recovered expression supports a declared endpoint only when high recovery is accompanied by tolerable edit burden, coherent marker context and stable downstream summaries.

**Table 1.**
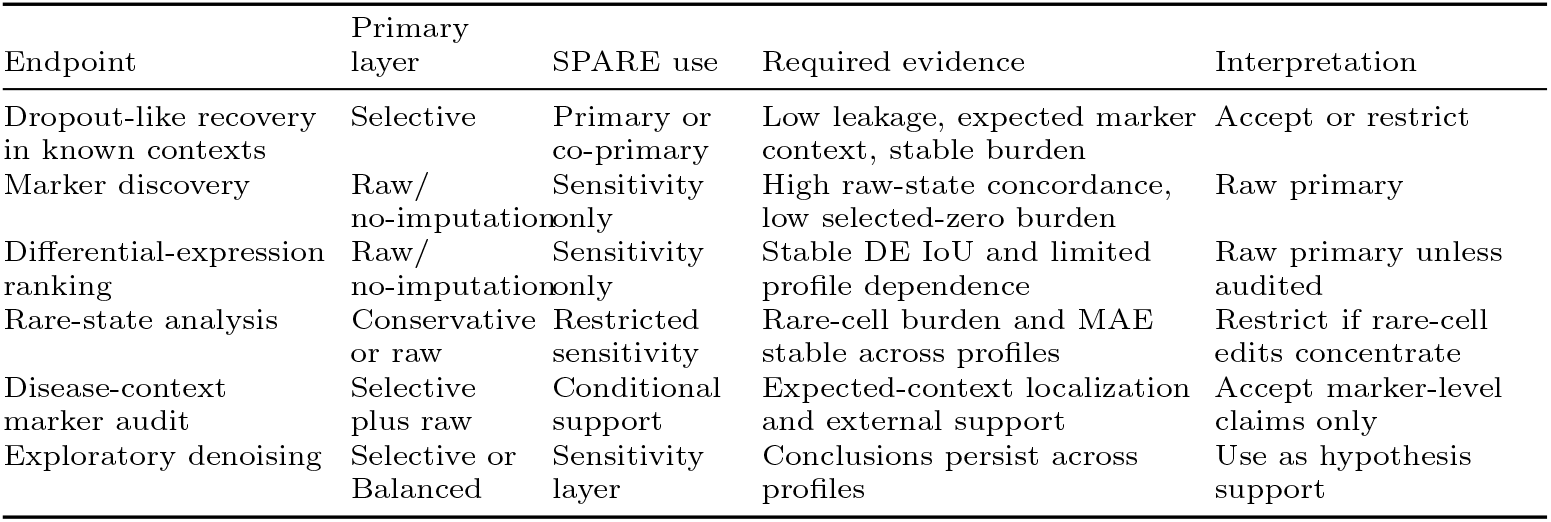
Endpoint-level reading guide for SPARE profiles and required audits.

### 2.2 A leakage-safe masking design separates recovery from answer-key contamination

When defining the benchmark for zero recovery we must avoid leaking held-out positives into the selection process. To ensure this separation we masked a subset of observed nonzero entries and split them into calibration and evaluation sets. The calibration entries were used to tune selector thresholds, while the valuation entries were reserved for scoring and were omitted from the selection mask. Under this protocol, global features such as, the principal component analysis (PCA), graph states and covariates, were recomputed from the masked baseline rather than carrying them over from the unmasked object. This ensured that the method received recovery credit only when its selection procedure independently identified an evaluation entry as recoverable.

To empirically evaluate its performance, we tested this framework on the PBMC 2700 (Zheng et al. 2017), Zeisel mouse brain (Zeisel et al. 2015), and Baron human pancreas datasets (Baron et al. 2016) Each benchmark utilized 2,000 highly variable genes (HVGs) spanning four mask fractions and three random seeds per fraction. Across these configurations we evaluated the no-imputation baseline, against the Selective SPARE reference profile and the Balanced/Conservative sensitivity profiles. The Selective profile reduced Root Mean Square Error (RMSE) relative to the no-imputation baseline by 41.6% in PBMC, 57.3% in Zeisel and 38.1% in Baron (see Table 2). These performance margins demonstrate that the selected-zero map successfully captures recoverable signal.

**Table 2.**
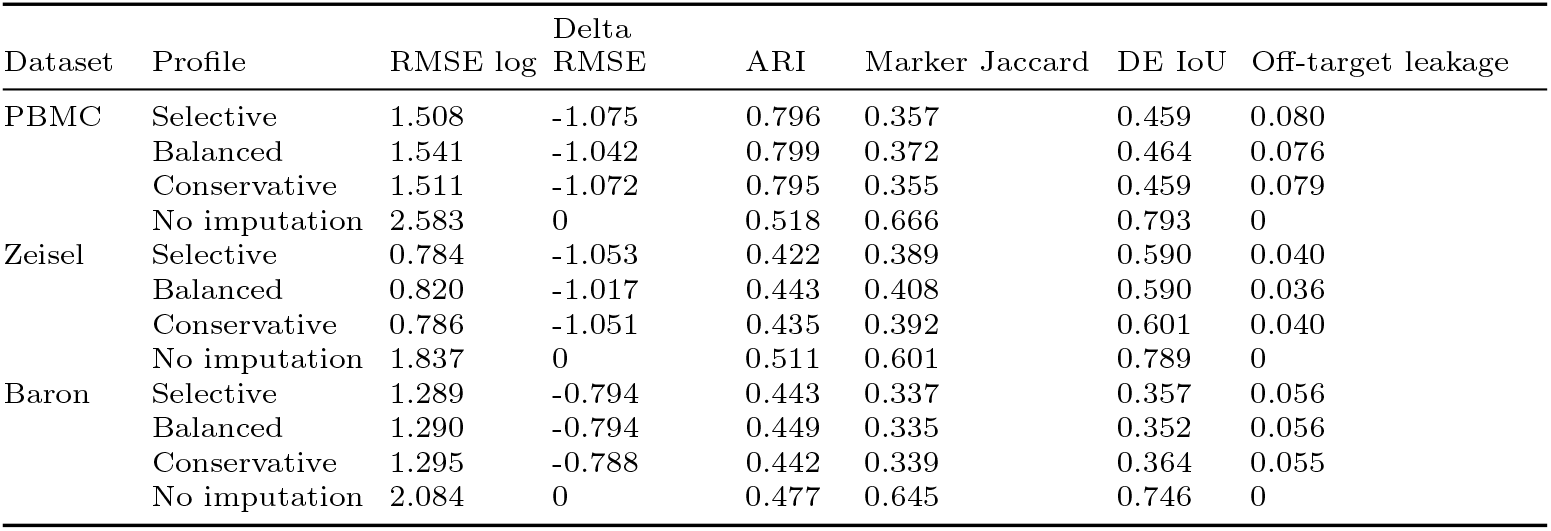
Compact 2000-HVG validation of no imputation, the Selective SPARE reference/recovery profile, and Balanced/Conservative operating-point sensitivity profiles across mask fractions 10/20/30/40% and three mask seeds per fraction. Values are means over the 12 matched run-level evaluations; standard deviations, burden columns, and per-mask summaries remain in the source-data tables and Supplementary Note 2. SPARE rows use the submitted optimized reconstruction backend.

Beyond global error reduction, evaluating the framework also requires monitoring how strongly raw marker and differential expression (DE) rankings were rewritten. These concordance metrics quantify descriptive behavioral shifts rather than intrinsic performance measures, as no external biological reference is utilized here. Internal rule-compliance leakage remained as zero for all SPARE configurations because rule-defined negative contexts are subjected to a strict hard-vetoed; additionally external marker panels establish the boundary for biological-zero risk The metrics reflected here (see Table 2) come from the clean calibration eval split suite and are not numerically interchangeable with their legacy comparator anchor.

However, high reconstruction fidelity alone is not the biological endpoint. SPARE also shifts marker and DE summaries away from the measured raw state. Modifying zero entries inherently modifies downstream rankings; therefore, this displacement is evaluated for biological defensibility rather than treated as a primary performance metric. For this reason, we interpret these metrics as a recovery-and-displacement audit rather than a one-dimensional performance ranking (see Table 2).

To navigate these distinct baseline metrics, SPARE is deployed across three operational profiles The Selective profile serves as the primary configuration optimized to capture recoverable biological signal. In contrast the Balanced and Conservative profiles acts as sensitivity variants for settings where prioritizing lower burden, rare-state preservation or raw-state concordance matters over maximal recovery. We therefore treat Selective as the main operating baseline, while Balanced/Conservative variants serve to test robustness.Thus, these profiles function as an integrated operational suite rather than separate, independent methods.

Internal rule-compliance functioned exactly as intended by maintaining zero leakage across the suite, this serves as a mechanical implementation check of the framework performance rather than a biological safety result as we intend. Thus we require a more rigorous safety boundary that instead leverages the marker contexts completely hidden from the selector or external to the internal rule table.

### 2.3 The intervention map exposes a recovery-cost frontier hidden by RMSE

In order to establish SPARE’s performance relative to the field, direct comparison against external imputers was conducted using ALRA and scVI as focused comparators under the same clean calibration/evaluation split (Linderman et al. 2022; Lopez et al. 2018). This clean rerun placed each method on a recovery-cost frontier rather than ranking methods by RMSE alone. This frontier combines masked observed-nonzero recovery, selected-entry burden, marker leakage and raw-state concordance.

In the focused clean rerun, scVI achieved the lowest RMSE across PBMC, Zeisel and Baron. ALRA also improved reconstruction, with particularly strong RMSE performance in PBMC and Baron. SPARE did not define the low-RMSE extreme of the frontier. Instead, it occupied a lower-burden and lower-leakage region while preserving native provenance for which observed zeros were selected before values were written (Table 3; Figure 3). This positioning represents an intentional architectural trade-off, where dense models can provide stronger reconstruction, but their completed matrices usually do not encode pre-write selected, rejected, untouched and vetoed states as first-class outputs.

**Table 3.** Focused clean comparator rerun under the same calibration eval split protocol used for the SPARE clean benchmark.

| Dataset | Method | Mask | RMSE | Recall | Edits (M) | Leakage | Marker J. | DE IoU |
| --- | --- | --- | --- | --- | --- | --- | --- | --- |
| PBMC | No imputation | None | 2.583 | 0 | 0 | 0 | 0.666 | 0.793 |
|  | SPARE Selective | Native | 1.508 | 0.499 | 0.386 | 0.080 | 0.357 | 0.459 |
|  | SPARE Balanced | Native | 1.541 | 0.484 | 0.373 | 0.076 | 0.372 | 0.464 |
|  | SPARE Conservative | Native | 1.511 | 0.497 | 0.383 | 0.079 | 0.355 | 0.459 |
|  | ALRA | Proxy | 0.988 | 0.962 | 3.297 | 0.208 | 0.143 | 0.256 |
|  | scVI | Proxy | 0.888 | 1 | 4.183 | 0.526 | 0.138 | 0.231 |
| Zeisel | No imputation | None | 1.837 | 0 | 0 | 0 | 0.601 | 0.789 |
|  | SPARE Selective | Native | 0.784 | 0.861 | 1.115 | 0.040 | 0.389 | 0.590 |
|  | SPARE Balanced | Native | 0.820 | 0.839 | 1.052 | 0.036 | 0.408 | 0.590 |
|  | SPARE Conservative | Native | 0.786 | 0.858 | 1.102 | 0.040 | 0.392 | 0.601 |
|  | ALRA | Proxy | 0.857 | 0.945 | 1.560 | 0.143 | 0.391 | 0.593 |
|  | scVI | Proxy | 0.528 | 1 | 2.698 | 0.660 | 0.369 | 0.539 |
| Baron | No imputation | None | 2.084 | 0 | 0 | 0 | 0.645 | 0.746 |
|  | SPARE Selective | Native | 1.289 | 0.637 | 3.301 | 0.056 | 0.337 | 0.357 |
|  | SPARE Balanced | Native | 1.290 | 0.637 | 3.302 | 0.056 | 0.335 | 0.352 |
|  | SPARE Conservative | Native | 1.295 | 0.632 | 3.239 | 0.055 | 0.339 | 0.364 |
|  | ALRA | Proxy | 0.935 | 0.951 | 6.813 | 0.170 | 0.245 | 0.387 |
|  | scVI | Proxy | 0.647 | 1 | 10.727 | 0.656 | 0.264 | 0.371 |

*Reported values represent means across four mask fractions, each calculated from three distinct seeds. None denotes the masked baseline without zero editing. Native means SPARE’s pre-write selected-zero mask. Proxy means a post-hoc positive-gain edited-zero audit of a completed matrix; ALRA and scVI do not expose a native selected-zero mask.*

**Figure. 3.**
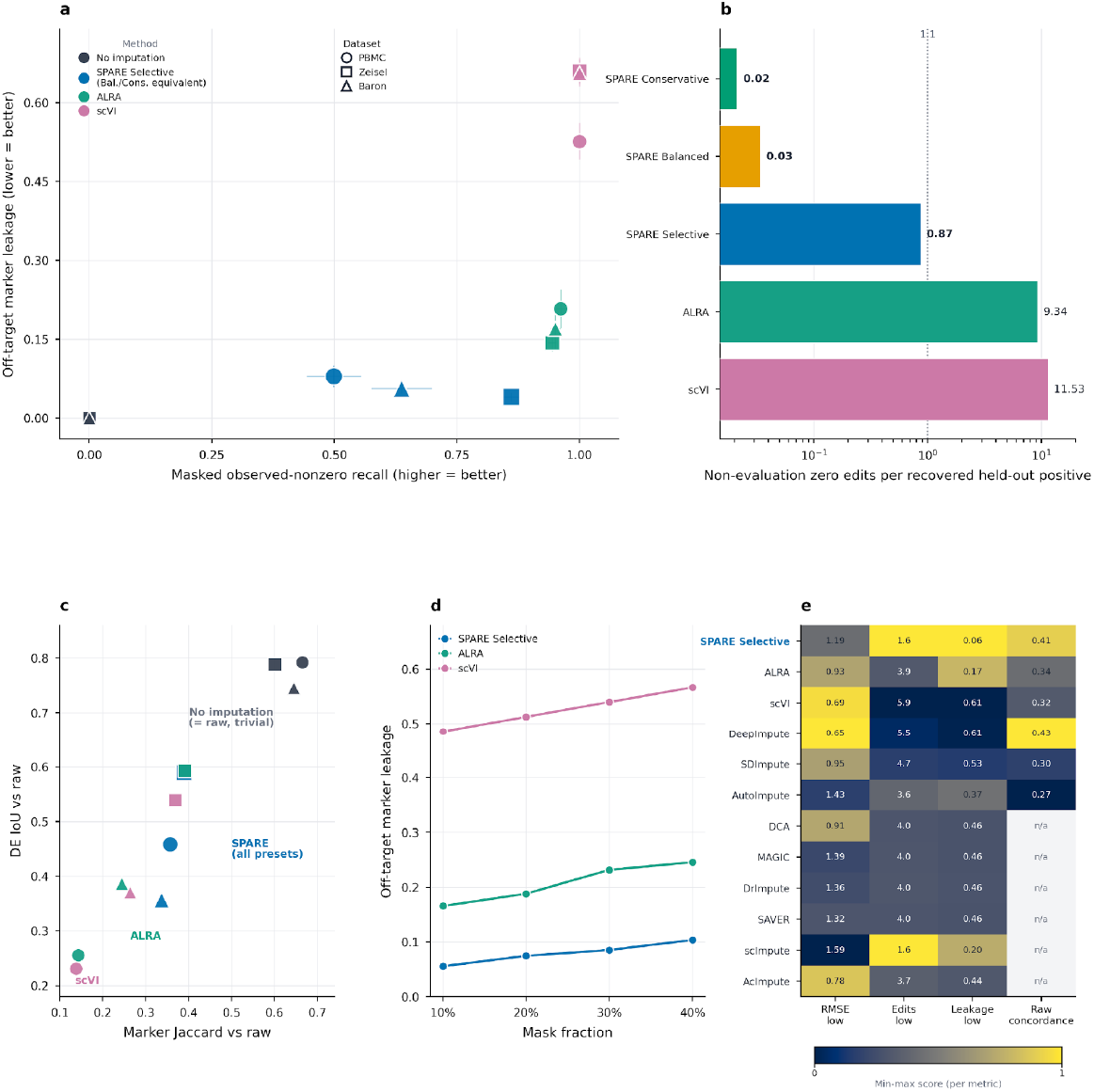
Comparator methods occupy different regions of the recovery-cost frontier. a, Masked observed-nonzero recall against off-target marker leakage for no imputation, SPARE Selective, ALRA, and scVI. SPARE Selective is shown as the representative SPARE point because Balanced and Conservative occupy the same operating region in this view. b, PBMC burden audit showing non-evaluation observed-zero edits per recovered held-out positive. c, Raw-state marker and differential-expression concordance; higher values indicate less movement from the measured-state analysis. d, PBMC mask-fraction sensitivity for off-target marker leakage. e, For each metric, raw values were converted to a favorable-direction score: lower-error, lower-burden and lower-leakage metrics were inverted, whereas recovery and concordance metrics were kept in their natural direction. Scores were then min-max scaled across the displayed comparator runs.

**Figure 4.**
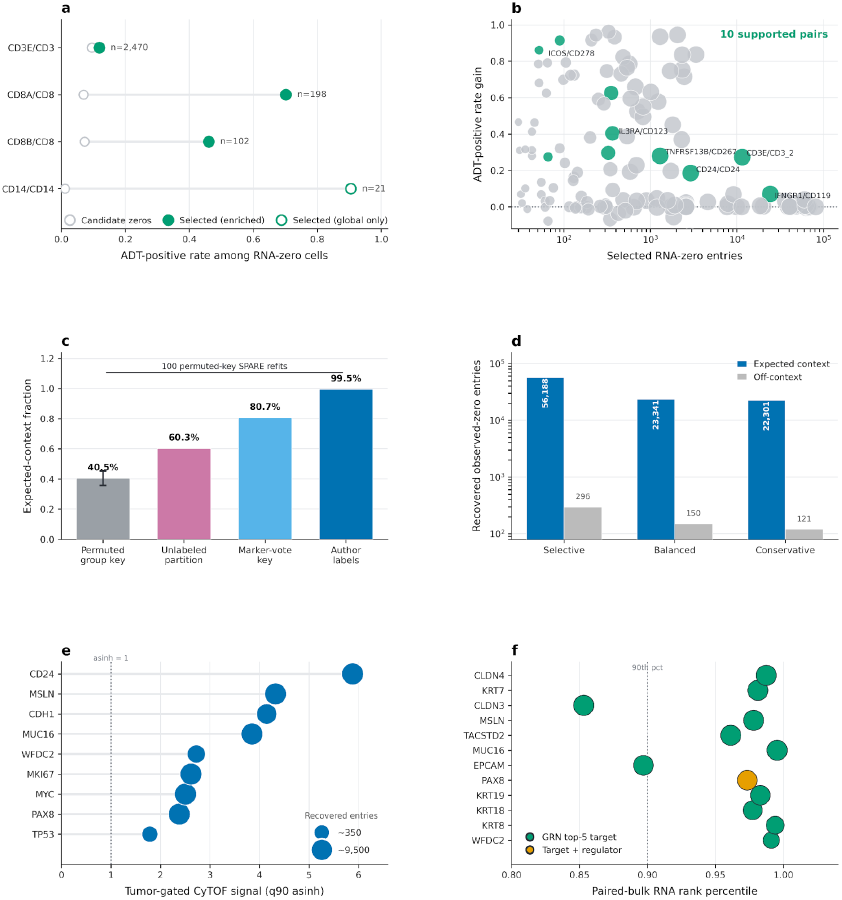
External protein and disease-context audits support endpoint-specific recovered signal. a, PBMC CITE-seq/TotalSeq-B audit comparing ADT positivity among candidate RNA-zero cells versus SPARE-selected RNA-zero cells, for predeclared immune surface markers; filled points denote selected zeros enriched within their partition, open points those supported only at the global level (n = selected RNA-zero entries per pair). b, Broad all-cells GSE164378 PBMC CITE-seq audit of RNA/ADT pairs (≥30 selected zeros); pairs with strict within-partition protein support, an ADT-positive rate gain ≥ 0.05 and ≥1.5-fold enrichment over candidate zeros are highlighted (10 supported pairs; 27 passed the broader RNA-partition-stratified global effect filter); point area scales with selected RNA-zero entries. c, HGSOC curated-marker audit showing the expected-context fraction of recovered zeros under increasingly informative grouping keys: 100 SPARE refits with deliberately permuted group keys (bar, mean; whisker, SD), unlabeled SPARE partitions, a marker-vote partition and author-defined cell types. The deposited source tables also include 10,000 no-refit context/label permutations as an additional null check, but those permutations are not the displayed bar. d, Expected-context versus off-context recovered observed-zero burden across SPARE presets (Selective, Balanced, Conservative; log scale). e, Tumor-gated CyTOF support for SPARE-lifted ovarian tumor markers (Gonzalez et al. 2018); the dashed line marks an asinh signal of 1 and point area scales with recovered entries (~350 to ~9,500). f, Paired-bulk RNA rank percentile for selected recovered tumor-marker genes; the dashed line marks the 90th percentile and point area scales with recovered entries. All twelve genes are GRNContext top-5 targets; PAX8 (orange) is additionally a local regulatory hub.

These results fundamentally alter how we interpret performance, precisely because a lower RMSE answer says that a model can reconstruct held-out observed values well. It does not determine whether the model’s natural-zero edits are biologically acceptable for marker discovery, differential expression or rare-state interpretation. SPARE makes this biological-use boundary measurable by reporting how many observed zeros were edited, where those edits occurred and whether they crossed marker-context boundaries.

The extended comparison panel for benchmarking includes diffusion, smoothing, newer methods and established legacy methods such as MAGIC (Dijk et al. 2018), and these are best interpreted as method-positioning context rather than as a secondary ranking. Several dense or broad-smoothing methods reached favorable reconstruction regions but carried higher edit burden, higher leakage or incomplete provenance. Splatter truth-aware simulations showed the same trade-off from another angle: broad recovery increased false-zero recall but also increased true-zero lift, whereas SPARE traded lower recall for higher selectivity. Ultimately, these trade-offs shift the paradigm from a universal method hierarchy toward an intentional frontier interpretation.

### 2.4 External marker, partition and rare-state audits define restricted-use boundaries

The main biological safety boundary comes from marker contexts that were not directly available to SPARE’s internal veto rules. We therefore evaluated selector-hidden curated-marker leakage, external-panel leakage, raw off-label expression and low-raw/high-lift contexts. These audits measure biological plausibility beyond the rules used by the selector itself.

PBMC and Zeisel showed moderate selector-hidden leakage, supporting use with external marker review. Baron was different. Selector-hidden leakage remained high across profiles, at approximately 41–43%, indicating that pancreatic endocrine and secretory contexts are more vulnerable to marker-context ambiguity (Table 4). In this setting, raw/no-imputation results should remain co-primary unless tissue-specific marker review, ambient/decontamination checks and low-raw/high-lift inspection support the edited contexts.

**Table 4.** Main safety-boundary summary across SPARE profiles.

| Dataset | Profile | Held-out leakage | External-panel leakage | Selector-hidden leakage | Recommended interpretation |
| --- | --- | --- | --- | --- | --- |
| PBMC | Selective | 0 | 0.021 | 0.130 | Use with external marker review. |
|  | Balanced | – | 0.019 | 0.119 | Use with external marker review. |
|  | Conservative | – | 0.021 | 0.129 | Use with external marker review. |
| Zeisel | Selective | 0.009 | 0.011 | 0.139 | Use with external marker review. |
|  | Balanced | – | 0.010 | 0.136 | Use with external marker review. |
|  | Conservative | – | 0.009 | 0.119 | Use with external marker review. |
| Baron | Selective | 0.026 | 0.018 | 0.431 | Keep raw/no-imputation co-primary. |
|  | Balanced | – | 0.018 | 0.431 | Keep raw/no-imputation co-primary. |
|  | Conservative | – | 0.018 | 0.414 | Keep raw/no-imputation co-primary. |

*Reported values represent the fraction of edited observed-zero entries, where 0 indicates zero leakage and an en-dash (–) denotes an unexecuted run. Internal rule-compliance leakage was 0 for all rows because rule-defined negative contexts are vetoed by construction. Generalized biological safety is judged from external off-rule columns, especially selector-hidden curated markers invisible to the selector safety table.*

The raw off-label audit explains why this result should not be read as a simple failure. Many nominally negative marker-label contexts already contained raw off-label expression before imputation. In Baron, most Selective off-label raw-zero lifts occurred in contexts where at least 20% of off-label cells already detected the marker, whereas only a small fraction occurred below 5% raw off-label detection. This pattern could reflect label granularity, shared endocrine programs, ambient signal, doublet-like mixtures, transitional biology or over-editing. The audit does not assign a cell-level mechanism. Instead, it identifies low-raw/high-lift contexts as the strongest restriction signal.

Partition quality provided a second boundary. SPARE is intentionally partition-aware, so noisy, coarse or mismatched partitions can change marker interpretation even when masked observed-nonzero recovery remains technically favorable. Rare-state audits added a related caution: Selective recovered more signal but increased rare-cell edit burden, whereas lower-burden profiles remained closer to raw in rare-state settings. The practical rule is endpoint-specific. Selective is appropriate for probing recoverable dropout-like signal; Balanced, Conservative or no imputation should be preferred when rare states, marker discovery or raw-state differential-expression ranking are primary.

### 2.5 Protein and disease-context audits support endpoint-specific recovered signal

We then evaluated independent protein support for selected RNA-zero edits. In a public PBMC CITE-seq/TotalSeq-B dataset, using the modality introduced for joint RNA and antibody-derived tag measurement (Stoeckius et al. 2017), SPARE was fit only on RNA features and scored against matched antibody-derived tag measurements. Selected RNA-zero entries for CD3E, CD8A, CD8B and CD14 were enriched for matched ADT-positive cells over burden-matched global nulls. A stricter within-partition analysis retained support for CD3E, CD8A and CD8B. CD14 remained protein-compatible in global and group-stratified scoring, but its selected zeros fell in small clusters that did not meet the within-partition contrast threshold.

A broader PBMC CITE-seq audit extended this analysis to the full 161,764-cell GSE164378/Seurat v4 3-prime multimodal PBMC reference. Among 136 RNA/ADT pairs with sufficient selected-zero support, 27 passed an RNA-partition-stratified protein-support filter, and 10 remained robust under the stricter within-partition effect filter. The strict set included T-cell, activation and immune-surface examples such as CD3E/CD3, IFNGR1/CD119, IL3RA/CD123, ICOS/CD278 and TNFRSF13B/CD267. LAMP1/CD107a was nominally enriched but failed the effect-size filter, illustrating how the audit rejects weak rows rather than converting all selected zeros into positive biological claims. These results provide partial cell-level protein corroboration for selected immune surface markers while preserving an important boundary: ADT positivity is independent protein evidence, not proof that every selected natural zero was a missed RNA molecule.

To ground the evaluation in a real disease context, we applied the framework to a high-grade serous ovarian carcinoma (HGSOC) dataset (Hippen et al. 2023). Rather than attempting unbiased marker discovery, this analysis evaluated the intervention map’s ability to distinguish context-plausible tumor-marker recovery from off-context marker lifting in a real disease dataset. In GSE217517, Selective SPARE was run on 54,137 annotated HGSOC cells and all 36,601 genes, with graph construction from 2000 HVGs. Global scRNA-bulk rank concordance changed only minimally after SPARE, so the biological interpretation rests on local marker context rather than global pseudobulk improvement.

A curated 41-gene ovarian tumor and microenvironment marker audit showed a partition-dependent localization gradient. Expected-context recovery was 60.27% when unlabeled SPARE partitions were used for fitting and author labels only for scoring, compared with 40.48% ± 4.79% across 100 SPARE refits with deliberately permuted group keys. A marker-vote broad group key increased localization to 80.66%, and author cell types set the expected-context ceiling at 99.48%. These controls show that HGSOC marker recovery depends on biologically meaningful partitions while retaining measurable support above unlabeled and permuted controls.

The raw-state denominator sharpened the claim. The 41-gene panel contained 1,778,074 observed-zero entries before SPARE, and SPARE recovered 56,484 of them. In expected marker contexts, raw detection was already 66.9%, and SPARE recovered 43.6% of the remaining expected-context observed zeros. In off-context rows, raw detection was 9.9%, and SPARE recovered only 0.018% of observed-zero entries. Thus, the HGSOC result supports context-localized recovery rather than broad marker spreading.

Orthogonal HGSOC layers supported restricted tumor-marker plausibility. Public tumor-gated CyTOF files measured protein for tumor markers overlapping SPARE-lifted genes, including MUC16, PAX8, WFDC2 and MSLN. GRNContext placed several lifted tumor markers in ovarian-cancer regulatory-network neighborhoods. Together with expected-context localization and raw same-context support, these layers support context-plausible dropout-like recovery for selected tumor markers. The same audit restricts or rejects stronger off-context immune/plasma claims such as GZMB, MZB1 and JCHAIN. HGSOC therefore provides the concrete accept/restrict/reject example for the intervention-map framework.

### 2.6 The same intervention map is exposed in AnnData workflows

The Python/AnnData implementation exposes the same audit objects used for the benchmark, protein-support and disease-context analyses. A standard SPARE imputation call writes the imputed layer together with the selected-zero mask, recoverability probabilities, per-cell burden, per-gene deltas, group-level summaries, marker-context tables and plotting-ready diagnostics. The software output is therefore not only a completed matrix; it is the intervention map plus the evidence needed to interpret it.

Figure 5 summarizes how these objects support endpoint-specific decisions. Users can identify which zero-rich genes were edited globally, where edits concentrate by group, whether off-label marker contexts already show raw expression, and whether low-raw contexts still receive selected-zero edits. These diagnostics can justify accepting an imputed layer for a declared endpoint, restricting it to a profile or gene set, rejecting low-raw/high-lift contexts, or escalating to decontamination, doublet filtering, reannotation or orthogonal validation.

**Figure 5.**
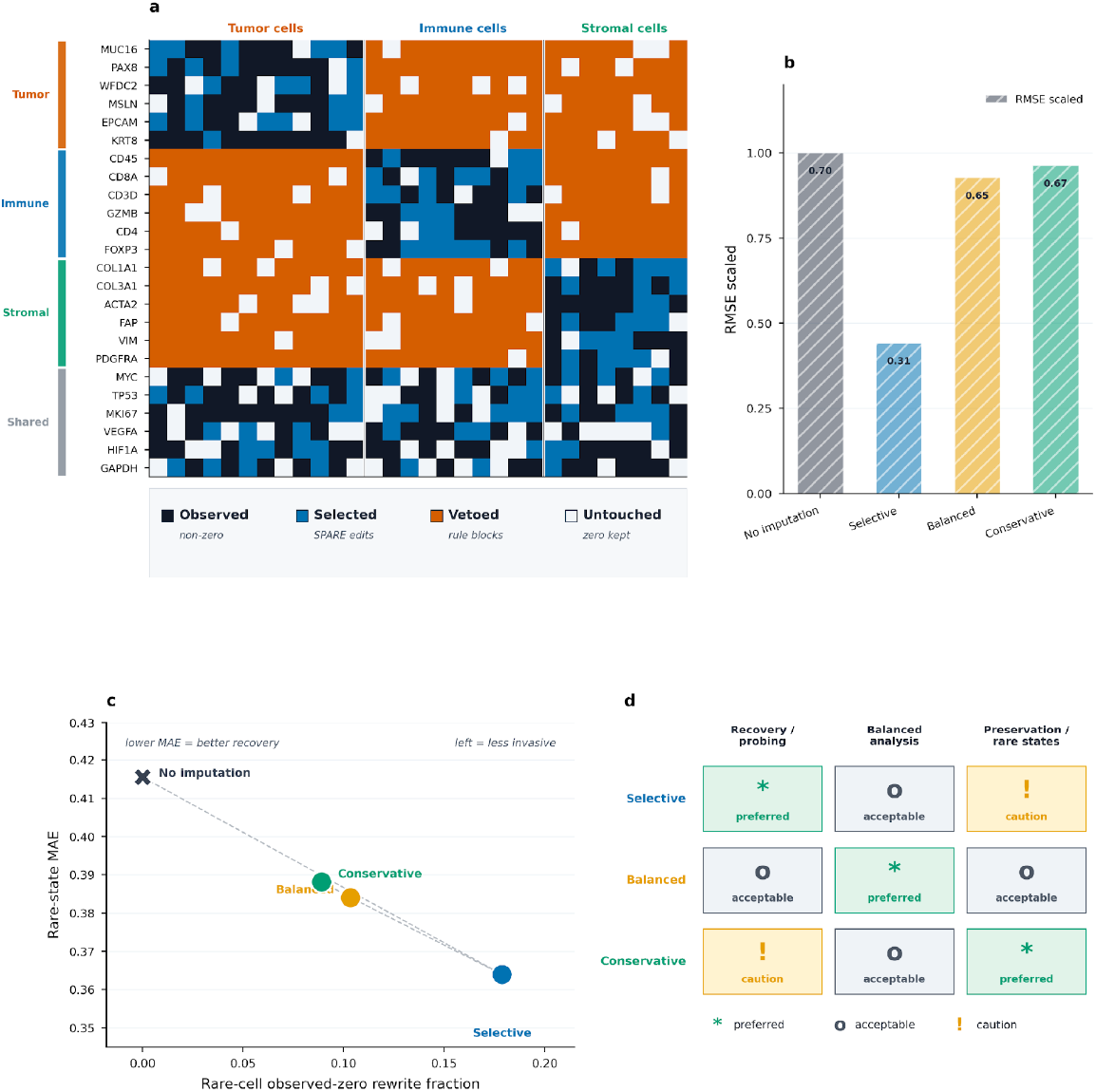
The AnnData implementation exposes the same intervention map used for biological decisions. a, SPARE stores measured nonzero entries, selected zeros, marker-vetoed zeros and untouched zeros as analysis objects rather than only writing a completed matrix; the schematic shows an illustrative 720-entry subset with 118 selected entries (16%), 311 vetoed entries (43%), 117 untouched zeros (16%) and 174 observed nonzero entries (24%). b, CellBench RMSE-scaled diagnostics. c, PBMC rare-state audit separating recovery from rare-cell edit burden. d, Endpoint guide mapping recovery, balanced analysis and preservation settings to Selective, Balanced and Conservative profiles.

Without this bundle, the workflow collapses back into ordinary matrix completion. A complete SPARE output therefore includes the imputed layer plus selected-zero provenance, burden summaries, marker-context audits and endpoint diagnostics. This design makes auditable zero recovery operational: recovered expression should be used only when the intervention that produced it remains visible.

## 3 Discussion

SPARE changes the unit of evidence in scRNA-seq imputation from the completed matrix to the edited observed zero. Its central contribution is the selected-zero intervention map, which records the observed zeros selected, left unchanged or marker-vetoed before reconstructed values enter downstream analysis. RMSE can show that a method recovers artificially masked observed nonzeros, but it cannot determine whether natural observed-zero edits are biologically defensible. By storing zero-selection, support and veto provenance before reconstruction, SPARE keeps recovery, burden, leakage and raw-state movement visible rather than absorbing them into a single completed matrix.

SPARE should not be interpreted as the first model to distinguish biological zeros from dropout-like zeros. That distinction has motivated zero-preserving, targeted, balanced and partition-aware imputation methods, including ALRA, scRecover, CPARI, SmartImpute, PbImpute and scZN (Linderman et al. 2022; Miao et al. 2025; Y. Zhang et al. 2025a, 2025b; Yao et al. 2025; Wu et al. 2026). SPARE’s contribution is instead one of provenance and endpoint interpretation. The selected-zero intervention map records the observed zeros selected, left unchanged or marker-vetoed before reconstructed values are written, allowing recovery to be interpreted together with edit burden, marker leakage, raw-state movement and external support.

This framing explains why SPARE should not be read as a universal low-RMSE replacement for dense imputers. In the clean comparator rerun, scVI achieved lower reconstruction error, and ALRA also performed strongly in several settings. Those results are important and should not be minimized. They show that dense models can be excellent reconstructors. SPARE addresses the complementary intervention end-point: which observed zeros were edited, which were left unchanged, which were vetoed, and whether those decisions remain defensible for a declared biological endpoint.

The selected-zero intervention map is most useful when imputation can affect biological interpretation. Marker discovery, differential expression, rare-state annotation and disease-context interpretation all depend on where non-detection is preserved or rewritten. In these settings, a completed matrix without provenance can obscure the difference between plausible dropout-like recovery and broad collateral editing. SPARE keeps that distinction visible by reporting recovery, burden, leakage and raw-state concordance together.

The audits also identify settings where raw data should remain primary. High selector-hidden leakage, low-raw/high-lift marker contexts, unstable rare states, weak partitions and unexplained downstream movement all argue against treating recovered expression as primary evidence. In those cases, SPARE should be used as a sensitivity layer or diagnostic tool rather than as the main analysis matrix. Conversely, when selected-zero burden is limited, marker context is coherent and external or same-context support is present, recovered expression can support a declared endpoint.

The biological evidence remains conditional. Masked observed nonzero entries provide direct labels for technical recovery, but natural observed zeros rarely have ground-truth labels. External marker panels, raw off-label patterns, partition perturbations, CITE-seq ADT measurements, HGSOC context localization, paired bulk, CyTOF and GRNContext provide different forms of support, but none proves that every edited natural zero was a missed RNA molecule. This limitation is not incidental; it is precisely why the intervention map is needed. It allows claims to be scaled to the evidence available.

The HGSOC and PBMC CITE-seq analyses illustrate this claim scaling. PBMC CITE-seq supports selected immune surface-marker recoveries at the protein level for a subset of markers, while rejecting others or restricting them to weaker evidence tiers. HGSOC supports context-localized tumor-marker recovery, particularly where raw same-context expression, CyTOF protein and regulatory-network evidence are concordant. It does not support unbiased marker discovery or broad off-context immune/plasma claims. These examples show how SPARE can support, restrict or reject recovered expression without pretending that imputation creates cell-level ground truth.

Several limitations follow directly from the design. SPARE depends on partition quality because it is intentionally partition-aware. Marker panels are incomplete and non-binary. Raw off-label expression can reflect label granularity, shared programs, ambient RNA, doublets, transitional states or over-editing. Profile choice was benchmark-driven rather than prospectively fixed for every possible endpoint. The present CITE-seq evidence is strongest for surface immune markers and does not validate non-surface genes or HGSOC tumor markers at cell-level resolution. These boundaries should remain explicit rather than hidden in supplementary material.

Future work should pre-register endpoint-specific profile choices, extend the clean comparator protocol to additional methods under matched resources, and test selected recovered signals with orthogonal assays such as spatial RNA, RNAscope/smFISH, disease-specific protein panels or matched multi-omic data. The practical rule is straightforward: imputed expression should be accepted for a biological endpoint only when the selected-zero intervention map, edit burden, marker context, raw-state concordance and downstream stability support that use.

## 4 Conclusions

SPARE reframes scRNA-seq imputation as auditable zero editing rather than hidden matrix completion. Its central object is the selected-zero intervention map, which records which observed zeros were selected, left unchanged or vetoed before downstream analysis.

Across clean benchmarks, comparator panels, protein-supported marker audits, disease-context analyses and AnnData examples, reconstruction error was informative but insufficient. The biological use of recovered expression depended on edit burden, marker context, raw-state movement, partition quality and external support.

In practice, SPARE should be interpreted endpoint by endpoint. Recovered expression can support dropout-like signal exploration when the intervention is limited and biologically coherent. It should remain a sensitivity layer when edits concentrate in low-support contexts, rare states or uncertain partitions. Raw/no-imputation analyses remain the measured-state reference unless the selected-zero audit supports a stronger claim.

## 5 Methods

### 5.1 Study Design And Scope

SPARE was evaluated as an auditable zero-recovery framework for scRNA-seq. The primary design combines leakage-safe masked observed-nonzero recovery, selected-zero burden quantification, marker-context leakage, raw-state concordance, comparator positioning and external biological audits. Imputation was therefore not treated as a single completed-matrix endpoint. Instead, the analyses determine when recovered expression can support specific downstream uses after the selected-zero intervention map has been inspected.

### 5.2 Data Scope

The primary validation datasets were PBMC 2700, Zeisel mouse brain, and Baron human pancreas. PBMC is a local project derivative of public 10x/Scanpy pbmc3k; Zeisel and Baron were exported from Bioconductor scRNAseq accessors and staged through repository scripts (Zheng et al. 2017; Zeisel et al. 2015; Baron et al. 2016). We then used external disease and direct-use datasets to test whether selected-zero edits remained interpretable outside the benchmark setting. These audits included SCP1288 renal cell carcinoma scRNA-seq with independent TCGA-KIRC bulk RNA-seq, GSE217517 paired HGSOC scRNA-seq/bulk RNA-seq (Hippen et al. 2023), independent HGSOC tumor-subset CyTOF FCS files (Gonzalez et al. 2018; Samusik 2018), the public 10x PBMC 10k CITE-seq/TotalSeq-B feature-barcode matrix, and the GSE164378/Seurat v4 multimodal PBMC CITE-seq reference for matched protein/RNA selected-zero audits (Hao et al. 2021). Dataset checksums, source manifests, and staging notes are maintained in the data availability and reproducibility files.

### 5.3 Preprocessing, Masking, And Partitions

Prepared count matrices were subset by stratified sampling when requested, normalized to a library-size log1p baseline, and restricted to 2000 HVGs for the main benchmark panels unless otherwise stated. The clean benchmark samples observed non-zero entries, masks them, and splits them evenly into calibration and evaluation sets. Calibration entries tune selector thresholds. Evaluation entries are reserved for scoring and count as recovered only if the method selects them independently. To prevent leakage, covariates, PCA, and graph state are recomputed from the masked baseline.

We distinguish three information categories. Label-free external comparators do not receive benchmark labels, although they may infer clusters internally. Label-aware external comparators receive benchmark labels directly; scimpute labeled is therefore reported separately as an information-aware comparator. SPARE is partition-aware: its configured group key contributes to local support, partition priors, recoverability scoring, and marker-safety contexts. Label perturbation, data-derived partition, marker-vote, and permuted group-key analyses test how sensitive the results are to that input.

### 5.4 SPARE Intervention Map, Zero Selection And Reconstruction

SPARE separates the decision to edit an observed zero from the reconstruction value assigned after that decision. The claim-bearing output is the selected-zero intervention map. This map distinguishes measured nonzero entries, selected observed zeros, marker-vetoed zeros and untouched observed zeros. Reconstruction is applied only after this selection and veto process, so a zero that fails selection remains unchanged even if a reconstruction engine could assign it a positive value.

Before entry-level scoring, SPARE builds a partition prior from the masked base-line and the configured group_key. Let *C*′ denote the masked count matrix, *B* = *log* (1 + *C*′_*normalized*_) the masked baseline, *c*_*i*_ the partition for cell *i, A* the row-normalized neighbor graph recomputed from *B*, and *Q*_*ij*_ the partition-prior expression for gene *j* in cell *i*. In the main benchmark path, each partition receives a gene-wise non-zero expression summary from cells in that partition; an optional cross-partition mode can mix this summary with similar partitions. Importantly, this prior is a contextual expectation, not an instruction to impute. It provides gene plausibility within a cell’s partition while preserving the distinction between detected expression, candidate recovery, and vetoed contexts.

The selector then scores each observed-zero entry by combining four sources of support. Negative-binomial evidence quantifies how surprising zero counts are given the gene’s empirical dispersion, the cell’s library size, and the partition mean. Local graph evidence measures detection or expression among neighboring cells in the masked baseline. Partition-prior evidence measures support from the cell’s group and related groups. Finally, gene-level recoverability summarizes prevalence, positive expression level, concentration in a top partition, and breadth of detected partitions. These gene statistics classify genes as rare, cluster-marker-like, or broad, and that class changes how much evidence is required before recovery is allowed.

For a cell-specific library factor *a*_*i*_, partition mean normalized count 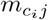, and empirical negative-binomial dispersion *θ*_*j*_, the NB support term is

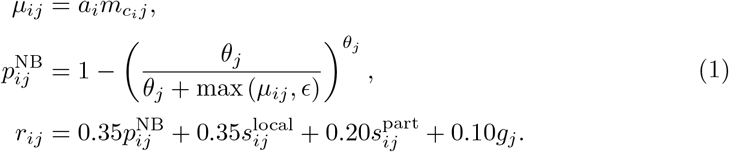

where *ϵ* is a small numerical floor, 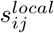 is local graph support, 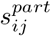 is partition-prior support, and *g*_*j*_ is gene-level recoverability. The reported profiles use the resolved configuration weights shown in this equation; profile-specific thresholds and ranker settings are listed in Supplementary Note 1.

Marker-safety rules are applied to partition-gene pairs before values are written. A gene can be treated as supported in its high-detection partition, neutral where evidence is ambiguous, or a negative marker in partitions where a rare or cluster-marker-like gene has little raw support. The reported profiles hard-veto rule-defined negative-marker contexts. For example, a tumor marker with local and partition support in tumor cells can remain eligible, whereas the same marker in an immune partition with negative-marker status receives zero recoverability probability.

Let *v*_*ij*_ ∈ *{*0, 1} denote the combined support and marker-safety gate. The base selected-zero probability is

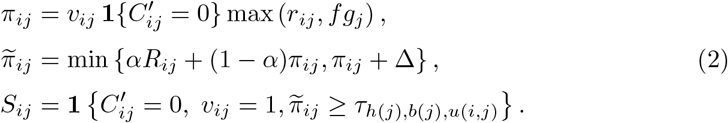

where *f* is the confidence floor and *R*_*ij*_ is the calibrated logistic-ranker probability trained on calibration-masked observed positives and burden-matched pseudo-negative natural zeros. The Selective profile uses *α* = 0.80 as the ranker/base-score blend and Δ = 0.18 as the maximum ranker-driven probability lift; Balanced and Conservative use stricter resolved values reported in Supplementary Note 1. Thresholds are chosen within gene-class, prevalence, and broad-context strata from calibration data; *S* is the selected-zero intervention map, *h*(*j*) is the gene class, *b*(*j*) is the prevalence stratum, and *u*(*i, j*) is the broad-gene context when used. Evaluation entries are not force-selected. Optional rescue extensions are allowed only through their own support gates and the same marker-safety constraints.

Profile names encode operating points rather than biological truth claims. Selective probes recoverable signal under audit; Balanced reduces editing pressure for endpoint sensitivity; and Conservative prioritizes raw-state preservation and marker-context caution. No imputation remains the measured-state reference in benchmark tables and downstream interpretation.

After the binary selected-entry mask is fixed, SPARE reconstructs values only for selected zeros. Genes are assigned to reconstruction sources according to their prevalence and partition specificity. Graph-supported genes use graph propagation anchored to observed entries; partition-supported genes blend graph information with the partition prior; and low-support genes use a low-rank reconstruction blended with contextual priors. Equation (3) describes the graph/partition-prior engine. Low-rank genes use the low-rank candidate source and then follow the same mask-only write rule. For each graph/partition-prior gene *j*, reconstruction is a weighted propagation problem on the masked-baseline graph. With *L* the graph Laplacian, SPARE estimates the cell vector 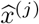 by minimizing

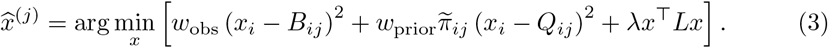

The first term anchors measured non-zero entries, the second anchors selected zeros to their contextual prior with recoverability-dependent weight, and the Laplacian term smooths values over graph neighbors. Under mask-only writing, observed non-zero entries remain anchored to the measured baseline and unselected zeros remain unchanged:

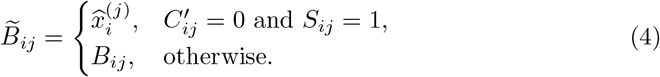

This write mode is why SPARE is evaluated as selective zero editing rather than blanket matrix completion.

### 5.5 Comparator Policy And Evaluation Endpoints

The focused clean external-comparator rerun uses the same calibration eval split protocol as Table 2 for no imputation, the three SPARE profiles, ALRA, and scVI. Broader comparator panels are retained as resource-matched method-positioning context across additional methods and newer-method stress tests, including DeepImpute, SDImpute, AutoImpute, DCA, MAGIC, DrImpute, SAVER, scImpute and AcImpute (Arisdakessian et al. 2019; Qi et al. 2021; Talwar et al. 2018; Eraslan et al. 2019; Dijk et al. 2018; Gong et al. 2018; Huang et al. 2018; Li and Li 2018; W. Zhang et al. 2025). Dense scVI output is audited with a post-hoc edited-zero proxy because scVI does not expose a native selected-zero mask. This proxy supports burden and leakage comparison, but it cannot recover pre-write support or veto provenance.

Endpoint classes are interpreted separately because they represent different evidence layers. Technical recovery endpoints include RMSE and masked observed-nonzero recall over evaluation masks. Intervention-scale endpoints include selected-entry burden, marker Jaccard, DE IoU, neighborhood preservation, correlation shifts, PAGA deltas, pseudobulk shifts, and signature shifts. Conditional biological endpoints include ARI, external curated-marker leakage, HGSOC expected-context recovery, paired bulk concordance, CyTOF overlap, CITE-seq ADT corroboration, GRNContext support, TCGA-KIRC aggregate projection, and marker-context nulls. Internal self-consistency endpoints include rule-compliance leakage and held-out rule-compliance leakage. We do not collapse these classes into a single score because doing so would obscure the distinction between recovery, intervention size, and biological plausibility.

### 5.6 Disease-Context And Software Audits

RCC direct-use, TCGA-KIRC projection, HGSOC paired-bulk, HGSOC marker-context, HGSOC CyTOF, CITE-seq ADT, and GRNContext analyses are observed-data plausibility audits rather than truth-labeled natural-zero benchmarks. The HGSOC 41-gene marker panel is explicitly post hoc and SPARE-informed, but it was curated from canonical ovarian tumor and microenvironment markers rather than selected by top SPARE metric rank. Matched, permuted, data-derived partition, marker-vote, and wrong-group-key controls estimate how much context localization exceeds alternative explanations within the same curated audit design.

For the PBMC CITE-seq audits, feature-barcode or Matrix Market inputs were split into gene-expression counts and ADT counts. SPARE was fit only on RNA features. The 10x PBMC 10k audit used external Cell Ranger graph-based RNA clusters as the partition key. The broader GSE164378 audit processed all 161,764 filtered cells in the 3-prime branch, retained 2,000 HVGs plus 202 ADT-mapped genes, and used RNA-only Leiden partitions as the primary group key; author celltype.l2 labels were retained only as sensitivity because the Seurat v4 annotations are multimodal. ADT-positive calls used a two-component mixture model after cell-wise CLR normalization and IgG/control-background subtraction. For each marker pair, observed RNA zeros were divided into candidate zeros and SPARE-selected zeros, and *P (ADT* ^+^|*RNA* = 0, *S* = 1) was compared with burden-matched global and partition-stratified null selections. The global null tests protein-compatible marker context; the partition-stratified null evaluates whether SPARE selects ADT-positive cells beyond the RNA-cluster prior. The GSE164378 broad-panel summary additionally reports an effect-size filter before interpreting nominally significant rows.

The spare-sc package implements SPARE for AnnData workflows. A typical call to sp.tl.impute writes the imputed layer, selected-zero mask, recoverability probabilities, burden summaries, gene/group tables, safety tables, thresholds, and diagnostics under AnnData layers, obs, var, and uns. The plotting API exposes these objects as endpoint diagnostics for global edited genes, group-resolved edited genes, marker lift, raw off-label expression, off-label lift, low-raw/high-lift contexts, label ambiguity, profile comparison, variance, burden, expression, delta, and UMAP summaries. These outputs guide endpoint-specific decisions; individual natural-zero validation requires independent evidence.

### 5.7 Reproducibility

The repository provides environment files, a Dockerfile, benchmark-suite configurations, figure builders, source-data tables, comparator wrappers, release-level checksums, and minimal rebuild commands for the clean validation, generated tables, disease-context audits, LaTeX export, and focused verification. Detailed command provenance, result manifests, supplementary source-data maps, and closure runbooks are included in the supplementary package so that the reported manuscript objects can be traced back to their generating commands.

ADT: antibody-derived tag; ARI: adjusted Rand index; CITE-seq: cellular indexing of transcriptomes and epitopes by sequencing; CyTOF: cytometry by time of flight; DE: differential expression; GRN: gene regulatory network; HGSOC: high-grade serous ovarian cancer; HVG: highly variable gene; NB: negative binomial; PAGA: partition-based graph abstraction; PBMC: peripheral blood mononuclear cell; RMSE: root mean squared error; scRNA-seq: single-cell RNA sequencing; TCGA: The Cancer Genome Atlas; ZINB: zero-inflated negative binomial.

## 5.8 Ethics approval and consent to participate

All analyses used previously published or public/reference single-cell and bulk RNA-seq datasets and generated no new human or animal participant data. No additional ethics approval was required for the secondary computational analyses reported here. Ethics approval and consent for the original datasets were handled by the original studies and data repositories.

## 5.9 Availability of data and materials

SPARE source code, benchmark manifests, figure builders, comparator wrappers, source-data tables and reproducibility documentation are available at https://github.com/ache1312/SPARE. The version described in this manuscript is archived at https://doi.org/10.5281/zenodo.20511804. Public datasets analyzed here are listed in the data manifest with accession identifiers, checksums, staging scripts and processing notes. Source-data tables and command provenance for all claim-bearing analyses are included with the archived release.

## 5.12 Authors’ contributions

SHG and AJMM conceived the project. SHG developed the software, implemented the benchmarking and audit pipeline, performed the computational analyses, generated the figures and tables, and drafted the manuscript. IPS, AHO, SR, ALl, and AJMM contributed to biological interpretation, validation strategy, and manuscript revision. ALl and AJMM supervised the work and acquired funding. All authors reviewed, edited, and approved the final manuscript.

## 5.13 Acknowledgements

The authors acknowledge the Centro Basal Ciencia & Vida and the NLHPC supercomputing infrastructure for institutional and computational support.

OpenAI ChatGPT were used for editorial language revision, manuscript-organization checks, and figure-design feedback during manuscript preparation. The authors reviewed, edited, fact-checked, and approved all manuscript text, code, analyses, citations, figures, and conclusions, and take full responsibility for the final content.

